# Single cell spatial analysis identifies regulators of brain tumor initiating cells

**DOI:** 10.1101/2022.09.13.507846

**Authors:** Reza Mirzaei, Charlotte D’Mello, Marina Liu, Ana Nikolic, Mehul Kumar, Frank Visser, Pinaki Bose, Marco Gallo, V. Wee Yong

**Author notes:** Correspondence to: V. Wee Yong, PhD, Professor, University of Calgary, 3330 Hospital Drive, Calgary, Alberta, T2N 4N1, Canada.

## Abstract

Glioblastomas (GBMs) are aggressive brain tumors with extensive intratumoral heterogeneity. Here, we used spatial transcriptomics and single-cell ATAC-seq to dissect the transcriptome of distinct anatomical regions of the tumor microenvironment. We identified numerous extracellular matrix (ECM) molecules including biglycan elevated in areas infiltrated with brain tumor-initiating cells (BTICs). Single-cell RNA sequencing showed that the ECM molecules were differentially expressed by cells including injury response versus developmental BTICs. Exogeneous biglycan or overexpression of biglycan resulted in a higher proliferation rate of BTICs, and this was associated mechanistically with LDL receptor-related protein 6 (LRP6) binding and activation of the Wnt/β-catenin pathway. Biglycan-overexpressing BTICs grew to a larger tumor mass when implanted intracranially in mice. This study points to the spatial heterogeneity of ECM molecules in the GBM microenvironment and suggests biglycan-LRP6 axis as a therapeutic target to curb GBM growth.

## Introduction

Glioblastoma (GBM) is the most common primary malignant brain tumor in adults ^1^. The current standard of treatment consists of maximal safe resection, followed by radiation and chemotherapy with oral alkylating agent temozolomide ^2^. Despite these interventions and new treatments such as tumor-treating fields, patients continue to have a median survival of fewer than 21 months ^3^. A significant factor contributing to the poor prognosis is the presence of brain tumor-initiating cells (BTICs) in the tumor microenvironment which drive initiation, growth, and recurrence of GBM ^4^. Thus, successful eradication of tumor will need the development of innovative treatment techniques that target BTICs in addition to the bulk of the tumor cells.

It has been increasingly appreciated that cellular heterogeneity within tumor (intratumoral heterogeneity) likely contributes to poor outcomes in GBM patients. Single-cell RNA-sequencing (scRNA-seq) studies have identified cell populations displaying multiple cellular programs including (i) neural progenitor cell-like (NPC-like), (ii) oligodendrocyte progenitor cell-like (OPC-like), (iii) astrocyte cell-like (AC-like), and (iv) mesenchymal-like (MES-like) state within the GBM microenvironment as described by several groups including our own ^5–8^. Although the cellular origin of these cellular programs is unknown, BTICs are assumed to be responsible for driving the intratumor heterogeneity ^9^. scRNA-seq has recently revealed that BTIC diversity is distributed along a transcriptional gradient spanning two major cellular programs: developmental and injury response signatures with differing degrees of proliferation and stemness capacity ^10^. However, because of loss of cell niche information after tissue dissociation in scRNA-seq, the underlying mechanisms driving diversity in BTIC phenotypes remain elusive.

Here, we applied spatial transcriptomics to a rodent model of GBM and used publicly accessible single-cell ATAC-seq along with single-cell RNA-seq data of human GBMs, and confocal protein imaging, to investigate the spatial heterogeneity of brain tumor microenvironment at a single-cell resolution. We found that different types of extracellular matrix (ECM) molecules were differentially expressed by cells including discrete types of BTICs. Among the upregulated ECMs, biglycan played an important role in promoting proliferation in BTICs, through interaction with LDL receptor-related protein 6 (LRP6) and downstream activation of the Wnt/β-catenin pathway.

## Results

### Spatial intratumoral heterogeneity in the GBM microenvironment

To understand factors that contribute to the growth of GBM cells, we performed spatial transcriptomics on brain tissue sections from mice intracranially transplanted with syngeneic mouse BT0309 BTIC line (Figure 1A). This line was generated from NPcis mice carrying mutations in *Nf1* and *Trp53* ^*11*^. The intracranial implantation of mBT0309 BTICs in mice recapitulated the key characteristics of a human World Health Organization grade 4 GBM ^12^. Brains were harvested on day 40 after tumor implantation following in vivo bioluminescence imaging (Figure 1B). The anatomic location of high proliferative regions was identified using hematoxylin and eosin (H&E) staining (Figure 1C). As a control, tissue from the same area of the brain was collected from healthy mice without tumor implantation (Figure 1C). To explore quality control (QC) metrics such as nCount_spatial (number of transcripts), nFeature-spatial (number of genes), percent_mito (mitochondria content) and percent_hb (hemoglobin content) on spatial transcriptomics data, the R package Seurat (v3) was used ^13^ (Supplementary Figure 1). Analysis of the spatial transcriptomics data was done using the 10x Genomics software Loupe Browser. K-means method was used to analyze clustering, and the number of clusters was set at eight (Figure 1D, E). To identify genes that are differentially expressed between the 8 clusters, we performed a differential gene expression analysis via the Locally Distinguishing method in the Loupe Browser (Supplementary Data 1). The heatmap in Figure 1F shows the top differentially expressed genes (DEGs) between the 8 clusters. Among the genes with differential expression, there were 10 genes encoding extracellular matrix (ECM) proteins including *Col9a1* (encoding collagen type IX, alpha 1), *Col9a2 (*encoding collagen type IX, alpha 2), *Col9a3* (encoding collagen type IX, alpha 3), *Col2a1 (*encoding Collagen type II, alpha 1), *Col11a1* (encoding collagen type XI, alpha 1), *Col11a2* (encoding collagen type XI, alpha 2), *Vcan* (encoding versican), *Fmod* (encoding fibromodulin), *Cspg4* (encoding chondroitin sulphate proteoglycan 4), and *Bgn (*encoding biglycan) that were expressed at higher levels in cluster 7 vs other clusters (Figure 1F). The higher expression of these ECM genes in cluster 7 is displayed by distributed stochastic neighbor embedding (t-SNE) plots (Figure 1G). Gene expression of these ECM genes was observed to be higher in tumor mice compared to normal brain (Figure 1H and Supplementary Figure 2A, B). In addition to cluster 7, there were also high transcript levels of *Bgn* and *Col9a3* in cluster 8 (Supplementary Figure 2C).

**Figure 1.**
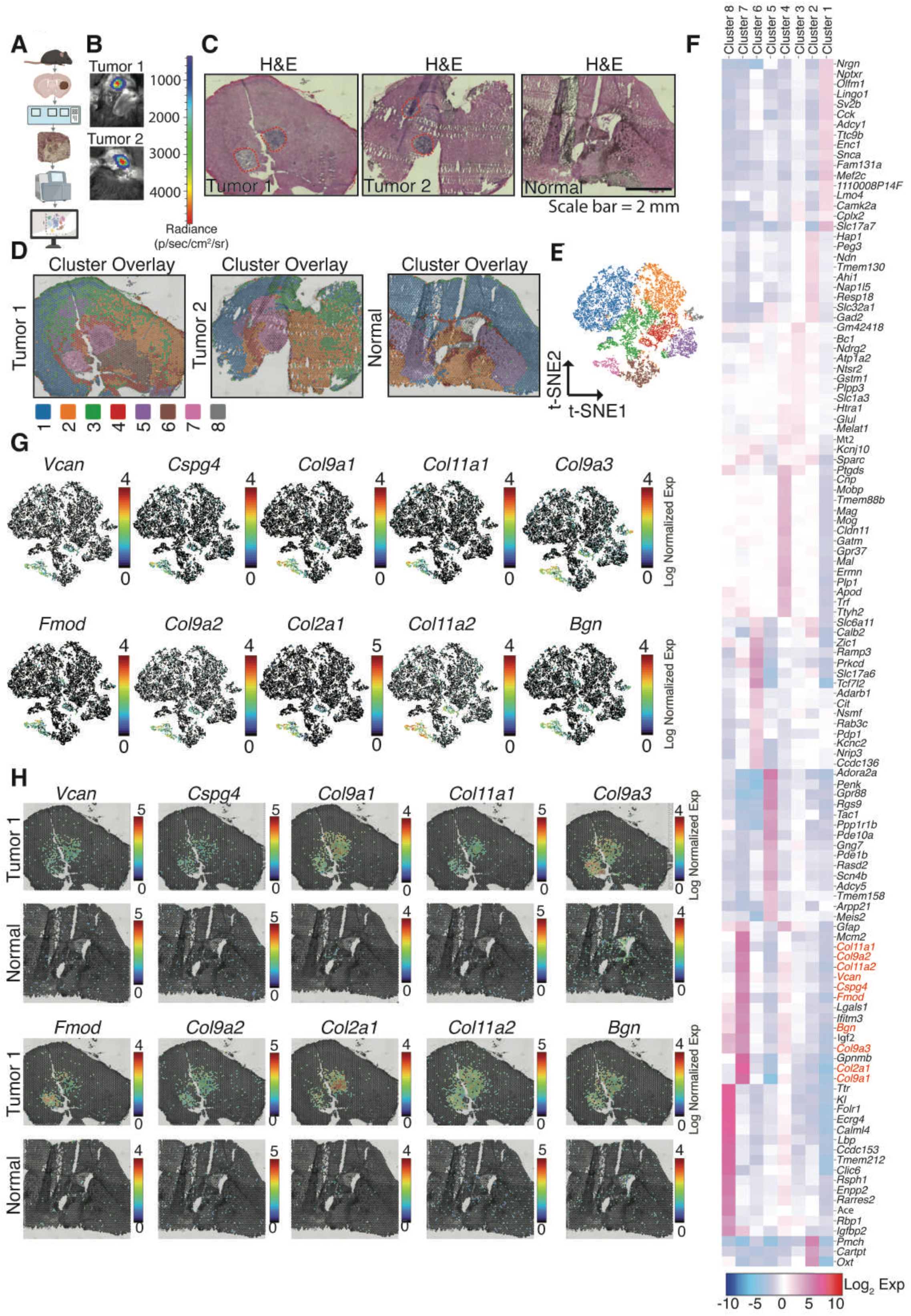
Spatial intratumoral heterogeneity in the GBM microenvironment. **A**. Schematic showing the spatially resolved transcriptomics experiment workflow. Forty days after intracranial implantation of syngeneic mouse BTICs, brain tissues were subjected to spatial transcriptomics and data analysis. **B**. Bioluminescence in vivo images of two mice 40 days after tumor implantation. **C**. Histological images (H&E) images of brain tissues obtained from two tumor mice and one healthy control without tumor injection. **D**. Cluster overlay images showing 8 clusters identified using K-means clustering. **E**. t-SNE plot of 8 clusters identified using K-means clustering. **F**. Heat map of top differentially expressed genes amongst the eight spatial clusters. **G**. t-SNE plots delineating distribution of the 10 ECM genes that were shown to be highly elevated in Clusters 7 and 8. **H**. Spatial distribution of the 10 ECM gene transcripts across tissue sections in tumor 1 and normal brain.

### Extracellular matrix molecules are differentially expressed by cells in the GBM microenvironment

Because the 10x Genomics spatial transcriptomics platform used here does not enable us to assess transcriptomic expression at the single cell level, we analyzed publicly available scRNA-seq data from 7 GBM patients ^10^ to dissect ECM gene expression on different cell populations. The R package Seurat (v3) ^13^ was used to perform quality control (QC), principal component analysis (PCA) and cluster analysis (Supplementary Figure 3A, B). Unsupervised clustering of all 44,712 cells from the 7 GBM patients, based on 2,000 variable genes and 15 significant principal components, delineated 17 clusters (Figure 2A). Lineage cell markers amongst the differentially expressed genes were used to identify the clusters. Clusters 0–2, 6, 9, 13, 14, and 17 were identified as microglia and monocyte-derived macrophages based on their expression of the following signature genes: *PTPRC* (CD45), *ITGAM, TMEM119*, and *CX3CR1* (Figure 2B and Supplementary Figure 3C). Clusters 11 and 16 were identified as T cells by their expression of *PTPRC* and *CD3E*. Clusters 3 and 15 were ascribed as oligodendrocytes based on their expression of *MAG* and *MOG*. We identified clusters 4, 5, 7, 8, 10 and 12 as tumor cells through their expression of putative tumor cell marker *EGFR*. Visualization of gene expression amongst the 17 clusters delineated that the 10 ECM genes described above were expressed at different levels amongst cells in the tumor microenvironment (Figure 2C). *COL11A2* and *COL9A3* were expressed in a subpopulation of oligodendrocytes and tumor cells. In contrast, *COL9A2* transcripts were detected in immune cells including microglia and monocyte-derived macrophages, oligodendrocytes and tumor cells. *COL11A1, COL9A1, FMOD, CSPG4* and *BGN* were mostly expressed by tumor cell clusters. Finally, we detected *VCAN* expression in a subpopulation of microglia and monocyte derived macrophages as well as tumor cells. *COL2A1* was not detected at appreciable levels by scRNA-seq.

**Figure 2.**
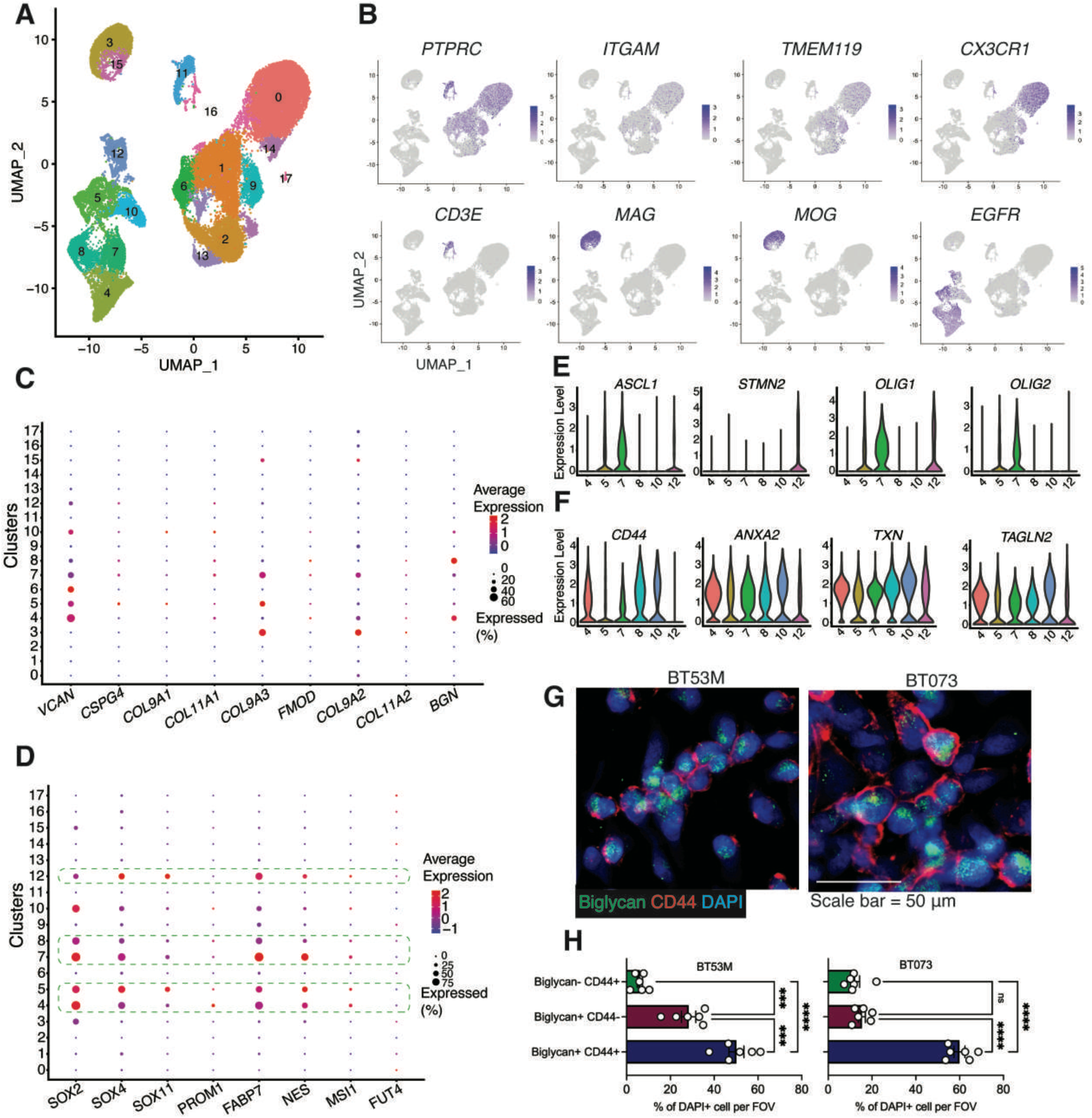
Differential expression of extracellular matrix molecules in cells within the GBM microenvironment. **A**. UMAP plot of 44,712 cells from 7 GBM patients. **B**. Feature UMAP plot showing the distribution of various genes used to define clusters representing microglia, monocyte-derived macrophages, T cells, oligodendrocytes and tumor cells in scRNAseq. **C**. Dot plot showing the expression of various ECM genes across the 17 cell clusters. The size of the dot corresponds to the percentage of cells expressing the gene in each cluster. The color represents the average gene expression level. **D**. Dot plot showing the expression of various stemness signature genes across the 17 cell clusters. Dashed lines show clusters with more expression of stemness transcriptome profile. **E, F**. Violin plots showing expression level of developmental (**E**) and injury response (**F**) signature genes. **G**. Representative widefield microscopy images of patient derived BTICs labeled with DAPI, biglycan and CD44. **H**. Bar graph comparing the percentage of cells expressing biglycan and/or CD44. Data are representative of two to three separate experiments. Means were compared between groups by one-way ANOVA: ***PLJ<LJ0.001, ****P < 0.0001. All data presented as the meanLJ±LJSEM (error bars).

To better understand the expression profile of the ECM molecules on BTICs, we first identified clusters 4, 5, 7, 8 and 12 as likely BTICs as they have higher expression of stemness genes including *SOX2, SOX4, SOX11, PROM1, FABP7, NES, MSI1* and *FUT4* (Figure 2D). A previous report has shown that the diversity of BTICs is distributed along a transcriptional gradient that spans two major cellular programs: developmental and injury response ^10^. We determined the expression of developmental versus injury response markers on tumor cell clusters. Clusters 5, 7 and 12 expressed higher levels of developmental markers such as *ASCL1, STMN2, OLIG1* and *OLIG2* (Figure 2E). In contrast, higher expression levels of injury response markers including *CD44, ANXA2, TXN* and *TAGLN2* were observed in clusters 4, 8 and 10 (Figure 2F). Single-cell expression analysis of the ECM genes showed that *COL11A1, COL11A2, COL9A1, COL9A2, FMOD, VCAN*, and *CSPG4* by both developmental and injury response tumor cells (Figure 2C). *COL9A3* was more expressed by developmental compared to injury response BTICs. Interestingly, *BGN* was either highly expressed or upregulated in tumor cells with more injury response than developmental phenotypes (Figure 2C).

Since genes associated with injury response tumor cells are associated with elevated expression of mesenchymal-related genes, we assessed the expression of *BGN* in different GBM subtypes through The Cancer Genome Atlas (TCGA) database. We observed significant upregulation of *BGN* in the mesenchymal subtype compared to classical and proneural subtypes (Supplementary Figure 4A). Examining the TCGA GBM Agilent-4502A and TCGA GBM RNA-seq databases confirmed positive significant correlations between *BGN* and *CD44* expression (r=0.34, p value=0.001; r=0.45, p value=0.001, respectively) (Supplementary Figure 4B). We also leveraged multiple public single-cell datasets to confirm a detailed characterization of cells expressing *BGN*. In this regard, we analyzed public scRNA-seq data from 21 adult GBM and 7 pediatric high-grade gliomas (HGG) resections encompassing 24,131 total cells ^5^. As previously described, most tumor cells in these 28 specimens clustered into cellular states according to multi-gene expression signatures ^5^. These categories were previously labeled as (i) NPC-like, (ii) OPC-like, (iii) AC-like, and (iv) MES-like states. We observed that there was a higher expression of *BGN* in MES-like cells than in other cellular states (Supplementary Figure 4C). Analysis of scRNA-seq data from 7 GBM patients also confirmed that *BGN* was either highly expressed or upregulated by clusters 4 and 8 exhibiting the injury response phenotype of BTICs (Supplementary Figure 4D).

To further corroborate the correlation between biglycan and mesenchymal phenotype of BTICs, we cultured patient derived BTICs and assessed the co-expression of biglycan and CD44 on BTICs (Figure 2G). Image analysis revealed that the percentage of biglycan and CD44 double-positive cells was significantly higher than the percentage cells expressing biglycan^+^ or CD44^+^ alone (Figure 2H).

### Higher gene activity of biglycan in BTICs within the GBM microenvironment

The spatial transcriptomics data from mouse GBMs revealed higher expression of stemness markers such as *Sox2, Sox4, Olig2* and *Fabp7* in areas with high expression of the ECM transcripts (Supplementary Figure 5A). There were higher expression levels of the stemness markers in malignant tissues compared to the normal brain (Supplementary Figure 5B). Furthermore, expression levels of stemness genes were upregulated in the spatial transcriptomic clusters 7 and 8 (Supplementary Figure 5C), suggesting these spatial clusters that were associated with higher *Bgn* expression exhibited more stemness signatures. To understand the chromatin accessibility of *BGN* gene, we examined a previously published dataset of single-cell chromatin accessibility in adult GBM ^8^. A significant increase in the gene activity of *BGN* was observed within the tumor compared to infiltrating non-neoplastic cells (Figure 3A). NMF decomposition of motif activity in the scATAC-seq data identified three cell types based on the stemness versus differentiation module: differentiated, intermediate, and stem cells (Figure 3B). *BGN* gene activity was shown to be significantly higher in stem cells compared to intermediate and differentiated cells (Figure 3C, D), confirming the higher expression levels of *BGN* in BTICs than other types of tumor cells.

**Figure 3.**
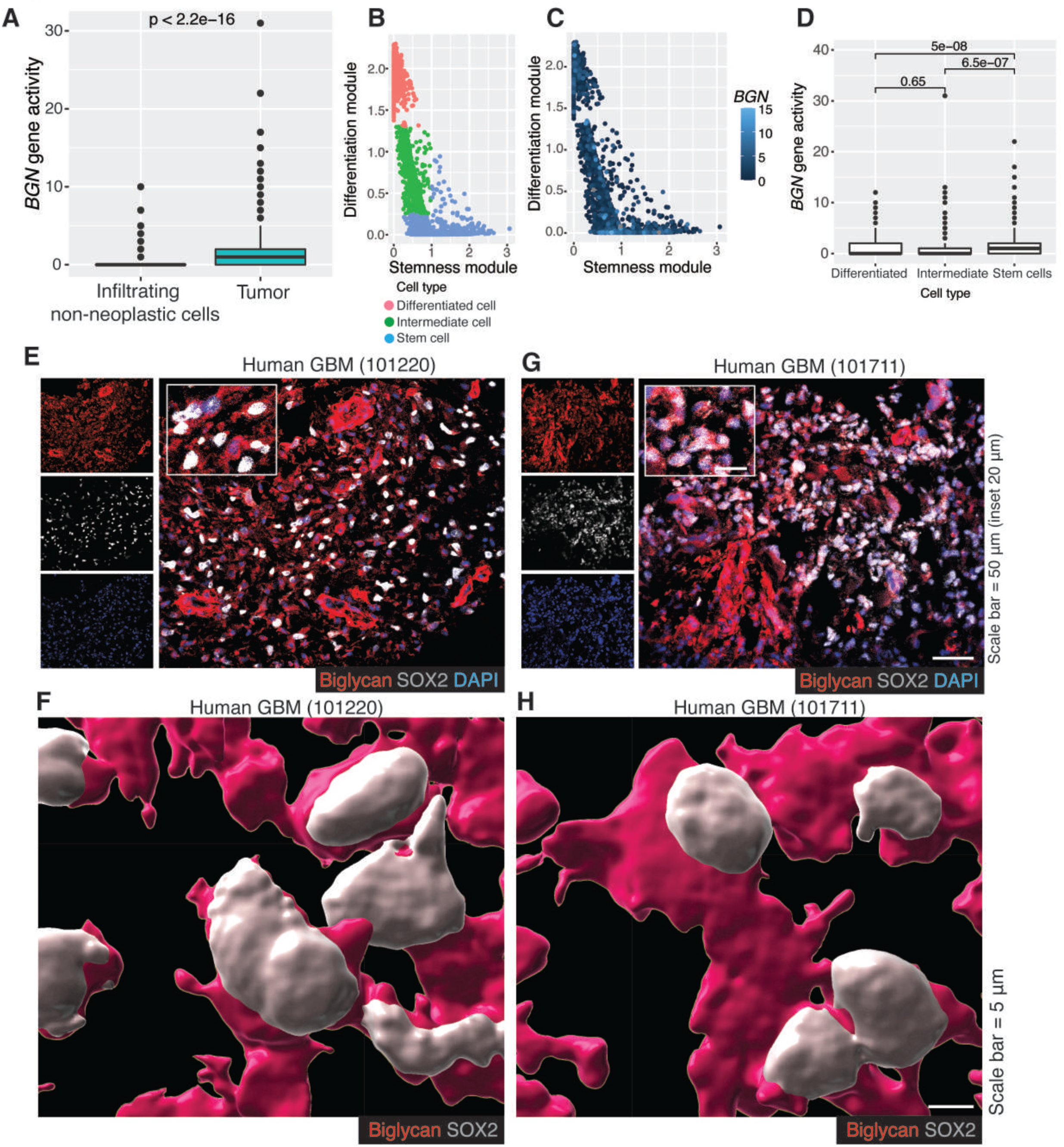
Higher gene activity of biglycan in BTICs within the GBM microenvironment. **A**. Analysis of scATAC-seq showed higher *BGN* activity in tumor compared to infiltrating non-neoplastic cells (P value <2.2e-16). **B**. Identification of three cell types in GBM based on differentiation and stemness modules in scATAC-seq. **C**. *BGN* activity in differentiation versus stemness modules. **D**. Bar graph comparing *BGN* activity in three cell types identified in scATAC-seq. **E, G**. Representative confocal staining assessing the proximity of biglycan and SOX2^+^ cells in human GBM tissues from two patients (101220, 101711). **F, H**. 3D reconstruction of images of SOX2, and biglycan in human GBM tissues.

Next, we evaluated biglycan expression in BTICs in human GBM specimens. While there is no uniform single marker for BTICs, SOX2 is widely used to identify BTICs within the GBM microenvironment ^14, 15^. Immunofluorescence confocal microscopy of human GBM specimens showed expression of biglycan by SOX2 positive cells (Figure 3E, G and Supplementary Figure 6A, B). Three-dimensional (3D) reconstruction of double labeling corroborated the presence of biglycan on SOX2 positive cells within the tumor microenvironment (Figure 3F, H and Supplementary Figure 6C, D). As controls, lower expression levels of biglycan and SOX2 were found in normal brains (Supplementary Figure 6 E-G). We were unable to detect staining signals after incubating human GBM tissues with secondary antibodies (Supplementary Figure 6H), ruling out secondary antibody reactivity with target markers. We also assessed biglycan expression on SOX2 positive cells in the mouse GBM model. Similarly, there was immunoreactivity of biglycan on SOX2 positive cells in the tumor microenvironment (Supplementary Figure 7A), which was corroborated by 3D reconstruction of double labeling (Supplementary Figure 7B). In contrast to tumor area, there was lower expression of biglycan and SOX2 in non-malignant area of mice implanted with tumor (Supplementary Figure 7C) and normal mouse brain (Supplementary Figure 7D).

### Biglycan is expressed in tumor areas with more proliferating cells

The spatial transcriptomics data demonstrated higher expression levels of proliferation markers such as *Mki67, Ccnd1, Ckap2l, Ccnd3*, and *Cdk4* in clusters 7 and 8, as seen in the violin plots in Figure 4A. In addition, spatial distribution of *BGN* transcripts across the tissue section showed higher expression of the proliferating markers in tumor areas with high levels of *BGN* transcripts (Figure 4B). Similarly, the scATAC-seq uncovered that cycling cells had significantly higher *BGN* activity than non-cycling cells (Figure 4C). Correlation analysis through the TCGA GBM Agilent-4502A and TCGA GBM RNA-seq databases confirmed positive significant correlations between *BGN* and *MKI67* expression (r=0.19, p value=0.001; r=0.29, p value=0.001 respectively) (Figure 4D). To validate the proximity of biglycan with proliferating BTICs at protein levels in the tumor microenvironment, we stained human GBM specimens for biglycan and the proliferative marker Ki67 as well as SOX2. Confocal images affirmed close proximity between proliferating BTICs and biglycan in the tumor areas (Figure 4E, F). Together, these data suggest that biglycan may be involved in promoting the proliferation of BTICs.

**Figure 4.**
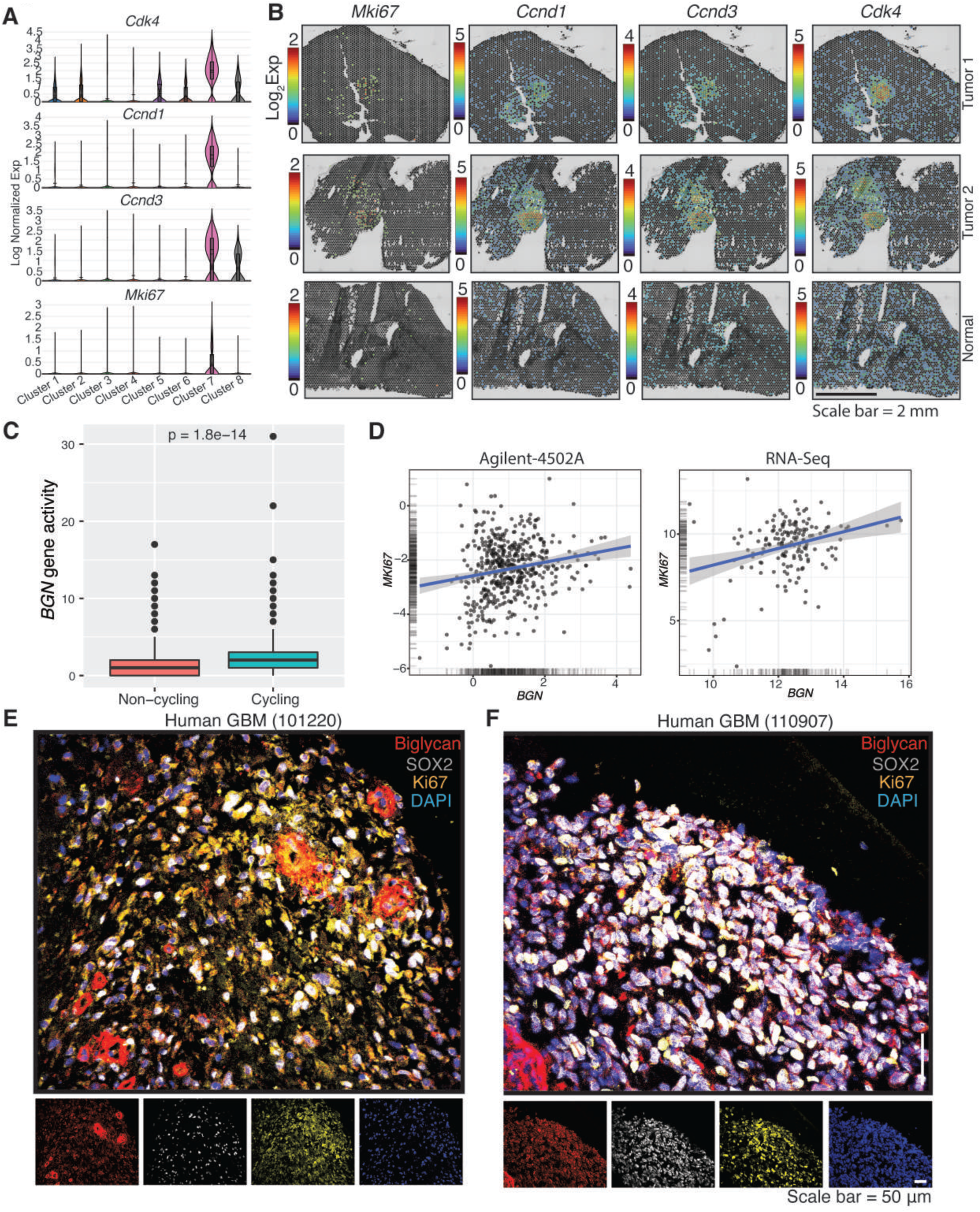
Expression of biglycan in tumor areas with more proliferating cells. **A**. Violin plots showing the expression levels of cell cycle genes amongst the 8 spatial clusters. **B**. Spatial distribution of cell cycle gene transcripts in tissue sections from tumor and normal brains. **C**. A bar graph comparing *BGN* activity in non-cycling versus cycling cells in human GBM tissues as determined by scATAC-seq analysis. **D**. Correlation between *BGN* and *MKI67* expression in the TCGA-GBM Agilent-4502A (left) and TCGA-GBM RNA-seq (right) database. **E, F**. Representative confocal images of human GBM specimens labeled with DAPI, SOX2, Ki67 and biglycan.

### Biglycan promotes proliferation of patient derived BTICs in culture

We next sought to understand the effects of recombinant biglycan on the growth of BTICs isolated from human GBM patients. After generation of BTICs from human GBM specimens, we treated cells with recombinant biglycan as shown in Figure 5A. Using 100 μm as the cutoff for sphere size, we found that exogenous biglycan increased sphere number in culture (Figure 5B, C). In addition, after 72 hours of treatment with recombinant biglycan, the ATP proliferation test revealed increased cell growth in BTICs (Supplementary Figure 8A). Flow cytometry EdU (ethynyl deoxyuridine) proliferation assay also showed that biglycan significantly elevated BTIC proliferation (Figure 5D, E). To confirm the proliferative effects of biglycan, we overexpressed *BGN* in patient derived BTICs. Immunofluorescence cell staining affirmed a significant upregulation of biglycan protein in *BGN* overexpressing (OE) cells compared to control (Figure 5F, G). Looking through clonogenic assays such as quantifying number of spheres showed that there was a significantly higher percentage of spheres in *BGN*-OE BTICs than control BTICs (Figure 5H). ATP proliferation assay revealed higher cell number in *BGN*-OE BTICs compared to control (Supplementary Figure 8B). Moreover, measuring sphere-forming capacity (SFC) of cells in the limited dilution assay indicated that *BGN*-OE BTICs produced significantly more spheres than controls (Figure 5I). Together, these data show that biglycan promotes proliferation of BTICs isolated from GBM patients.

**Figure 5.**
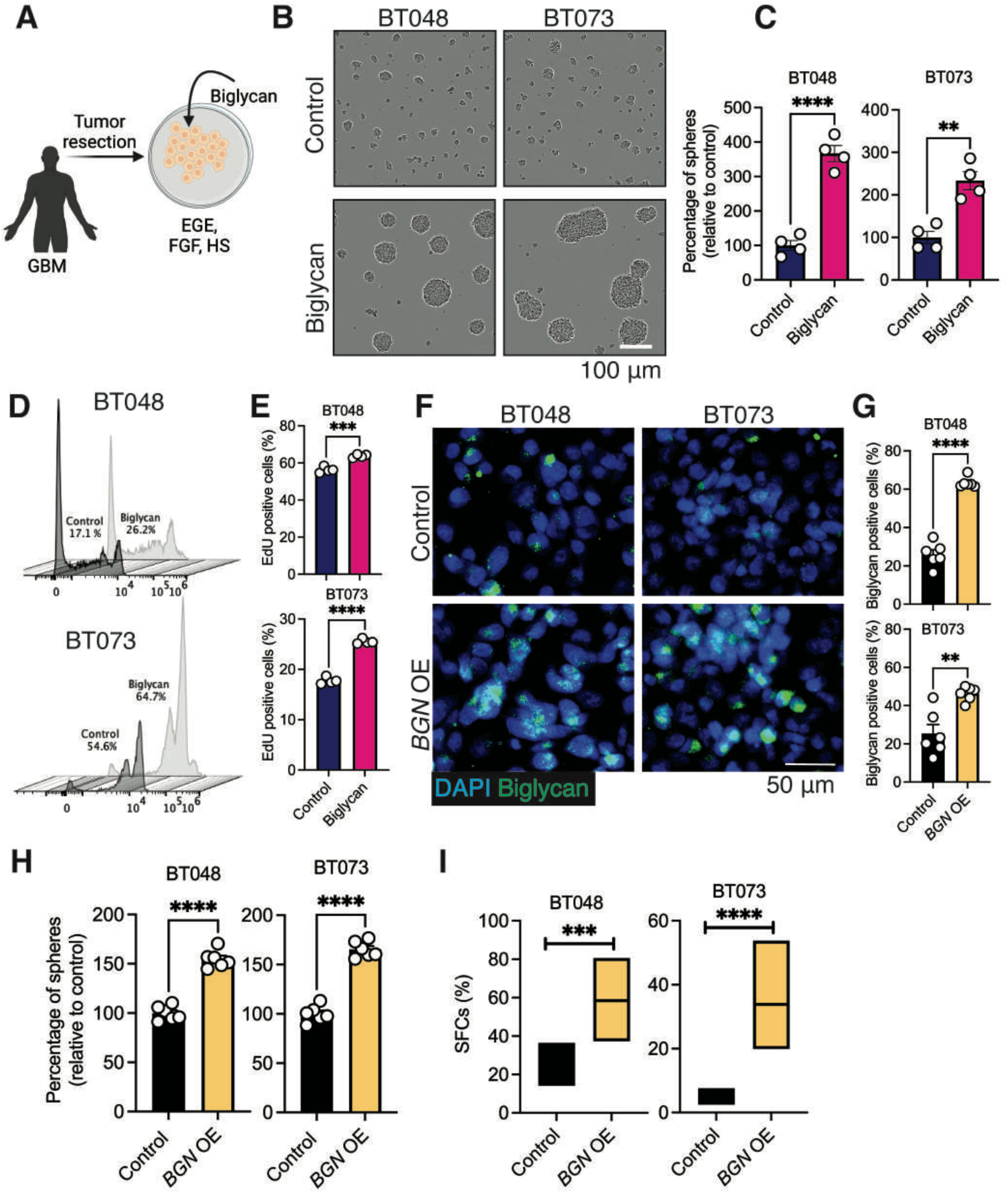
Biglycan promotes proliferation of patient derived BTICs in culture. **A**. Schematic showing the procedure of BTIC generation from patients with GBM and culture with recombinant biglycan. **B**. Representative bright-field microscopy images of 48-72-hour outcomes of tumor spheres. **C**. Quantification of tumor spheres of two human BTIC lines. Four fields of view (FOVs) from four cell culture wells were used to quantify number of spheres using 100 μm as the cutoff. **D, E**. Representative stagger offset plots (**D**) and bar plots (**E**) of the proliferation of human BTIC lines measured by flow cytometry of EdU proliferation assay after 24 hours of treatment. **F**. Representative widefield microscopy images of patient derived BTIC lines (BT048 and BT073) overexpressing *BGN* compared to control. Cells were labeled with DAPI and biglycan. **G**. Bar graph comparing the percentage of cells expressing biglycan between biglycan overexpressing cells compared to control. **H**. Bar graph comparing the percentage of spheres in *BGN* overexpressing BTICs versus control. **I**. Sphere forming capacity (SFC) of BTICs overexpressing *BGN* compared to control. Data are representative of two to three separate experiments. Means were compared between groups by unpaired (two-tailed) t test: **P < 0.01, ***P < 0.001, and ****P < 0.0001. All data presented as the meanLJ±LJSEM (error bars).

### Biglycan activates Wnt/β-catenin through interaction with LRP6 on BTICs

Based on the literature, there are several receptors that can bind to biglycan, including TLR2 and 4, P2X purinoceptor (P2RX) 2 and 4, and low-density lipoprotein receptor-related protein 6 (LRP6) ^16–18^. We assessed the mRNA expression of these receptors in two human BTIC lines and found higher expression of *LRP6* than other receptors (Figure 6A). In addition, scRNA-seq analysis of 7 GBM patients showed that compared to other receptors there were higher expression levels of *LRP6* on clusters 4, 5, 7, 8, 10 and 12 which correspond to tumor cells (Figure 6B). Spatial transcriptomics of mouse GBM confirmed anatomical expression of *Lrp6* in tumor area corresponding to cluster 8 (Figure 6C, D). Based on the role of biglycan in BTIC proliferation, we analyzed the TCGA GBM Agilent-4502A and TCGA GBM RNA-seq datasets and found a positive association between *MKI67* and *LRP6* (r=0.34 and r=0.39, respectively) (Figure 6E). Co-labeling of human GBM tissues confirmed expression of LRP6 on SOX2 positive cells which are in proximity to biglycan (Figure 6F). Together, these findings point to LRP6 as a possible biglycan receptor on BTICs and that blocking this receptor may reduce biglycan proliferative effects. To test this, we treated *BGN*-OE BTICs with monoclonal antibodies against LRP6. After 48 hours, *BGN*-OE cells treated with blocking antibodies had significantly reduced proliferation levels compared to cells incubated with isotype control antibodies (Figure 6G).

**Figure 6.**
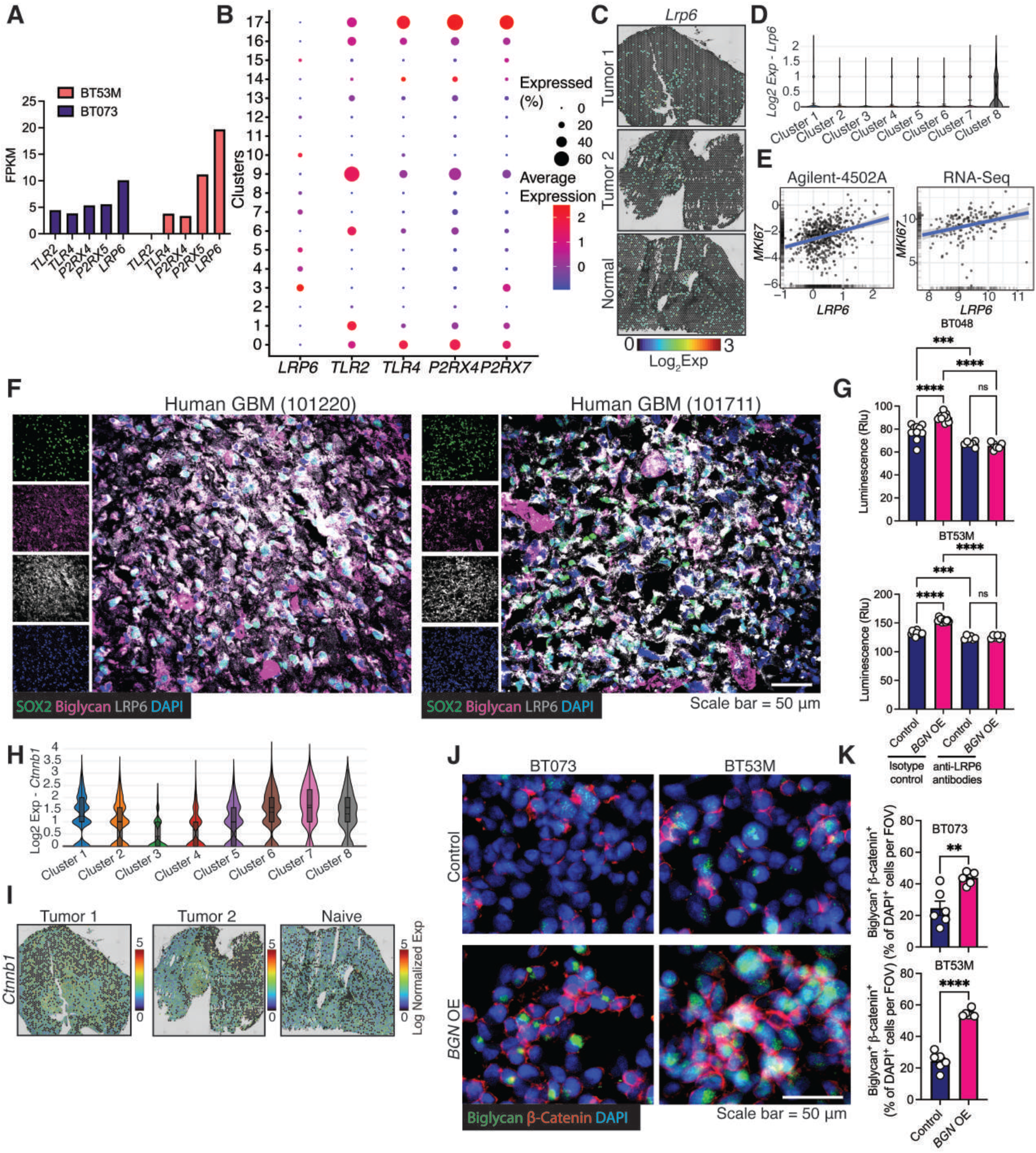
Biglycan activates the Wnt/β-catenin through interaction with LRP6 on BTICs. **A**. Expression levels of biglycan receptors in two human BTIC lines determined via bulk RNA sequencing. Fragments per kilobase of exon per million mapped fragments (FPKM). **B**. Dot plot showing the expression of various biglycan receptors across 17 single-cell cell clusters. The size of the dot corresponds to the percentage of cells expressing the gene in each cluster. The color represents the average gene expression level. **C**. Spatial distribution of *Lrp6* transcripts across tissue section in tumor and normal mouse brains. **D**. Violin plot displaying expression levels of *Lrp6* gene transcripts amongst the 8 spatial clusters. **E**. Correlation between *LRP6* and *MKI67* expression in the TCGA-GBM Agilent-4502A (left) and TCGA-GBM RNA-seq (right) database. **F**. Representative confocal microscopy images of human GBM specimens. Tissue was labeled with DAPI, SOX2, biglycan and LRP6. **G**. ATP proliferation assay of human BTICs overexpressing *BGN* after 48-72 hours treatment with LRP6 blocking antibody. **H**. Violin plot displaying expression levels of *Ctnnb1* transcripts (encoding β-catenin) amongst the 8 spatial clusters. **I**. Spatial distribution of *Ctnnb1* transcripts across tissue sections in tumor and normal mouse brains. **J**. Representative widefield microscopy images of patient derived BTICs overexpressing biglycan and control. Cells were labeled with DAPI, biglycan and β-catenin after overnight incubation. **K**. Bar graph comparing the percentage of cells expressing biglycan and β-catenin. Data are representative of two to three separate experiments. Means were compared to respective control by unpaired (two-tailed) t test when comparing two groups. For more than two groups, one-way analysis of variance (ANOVA) with Tukey’s post hoc was used: **P < 0.01, ***P < 0.001, and ****P < 0.0001. All data presented as the meanLJ±LJSEM (error bars).

Previous studies demonstrated LRP6 as an upstream driver of the Wnt/β-catenin signaling pathway ^19^. Spatial transcriptomic analysis of the tumor microenvironment showed high expression of *Ctnnb1* gene in clusters 7 and 8 (Figure 6H). Indeed, *Ctnnb1* was upregulated by cells infiltrated into tumor areas when compared to normal brain, which showed a diffuse expression at lower levels (Figure 6I). These observations suggested that activation of Wnt/β-catenin may be downstream of the LRP6-biglycan axis. To test this, we quantified β-catenin expression in *BGN*-OE cells compared to control BTICs (Figure 6J). After overnight culture, there were significantly higher expression levels of β-catenin in *BGN*-OE BTIC than control (Figure 6K).

### Biglycan accelerates in vivo tumor growth in the GBM animal model

To address in vivo growth benefits of biglycan on tumor growth, we implanted syngeneic mouse BT0309 with stable up-regulated *Bgn* into the striatum of immunocompetent C57BL/6 mice. By day 45, we found that mice with up-regulated *Bgn* BTICs had significantly higher tumor burden compared to control mice as observed by in vivo bioluminescence imaging (Figure 7A, B). Staining for Ki67 as an index of cell proliferation showed significantly higher immunoreactivity in mice implanted with *Bgn*–overexpressing cells compared to control (Figure 7C, D). To determine if in vivo biglycan overexpression led to activation of the Wnt/β-catenin pathway, we stained tumor tissues for β-catenin. Forty-five days after tumor implantation, there were higher expression levels of β-catenin in the tumor of mice implanted with *Bgn* overexpressing BTICs than control (Figure 7E, F). Staining for CD44 as an index of mesenchymal state also showed higher immunoreactivity in SOX2-positive BTICs in mice implanted with *Bgn* overexpressing cells compared to control (Figure 7G, H).

**Figure 7.**
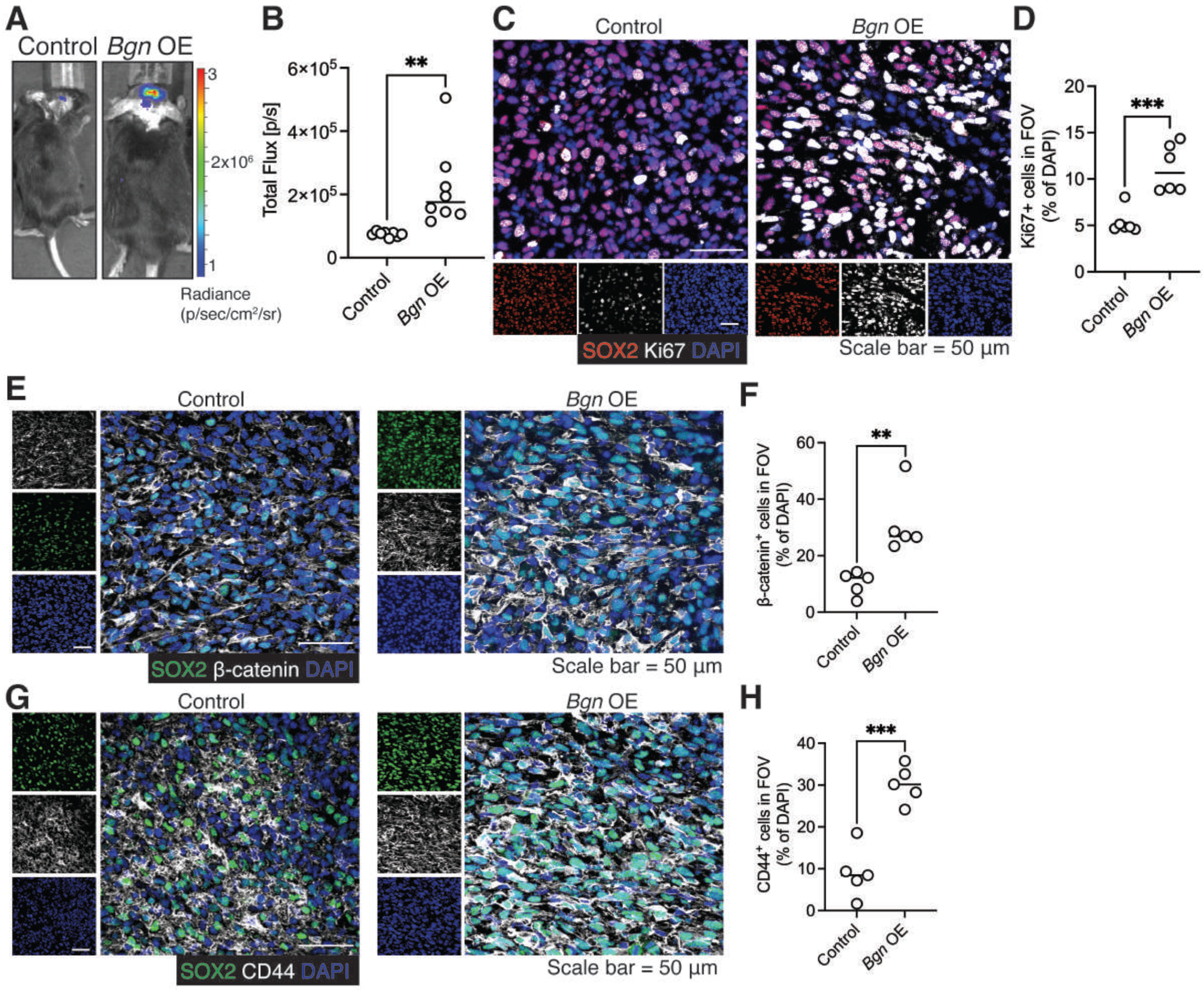
Biglycan accelerates in vivo tumor growth in the GBM animal model. **A**. Representative images of in vivo bioluminescence monitoring of tumor growth in mice implanted with *Bgn* overexpressing mouse BTICs compared to than in mice grafted with control cells. **B**. Graph comparing total Flux signal in mice implanted with *Bgn* overexpressing BTICs versus control. **C**. Representative confocal microscopy images of mouse GBM tissues implanted with *Bgn* overexpressing BTICs versus control. Tissue was labeled with DAPI, SOX2 and Ki67. **D**. Graph comparing the percentage of Ki67 positive cells between mice grafted with *Bgn* overexpressing BTICs versus control. **E**. Representative confocal microscopy images of mouse GBM tissues implanted with *Bgn* overexpressing BTICs versus control. Tissue was labeled with DAPI, SOX2 and β-catenin. **F**. Graph comparing the percentage of β-catenin positive cells between mice grafted with *Bgn* overexpressing BTICs versus control. **G**. Representative confocal microscopy images of mouse GBM tissues implanted with *Bgn* overexpressing BTICs versus control. Tissue was labeled with DAPI, SOX2 and CD44. **H**. Graph comparing the percentage of CD44 positive cells between mice implanted with *Bgn* overexpressing BTICs versus control. Data are representative of two to three separate experiments. Means were compared between groups by unpaired (two-tailed) t test: **P < 0.01, ***P < 0.001.

## Discussion

Here, we used spatially resolved and single-cell and single-cell ATAC transcriptomic approaches to dissect BTIC heterogeneity and reveal key regulators of their diversity in the GBM microenvironment. Spatial transcriptomics and scRNA-seq analyses in mouse and human GBM highlighted the heterogeneity of BTICs and identified several ECM molecules which are differentially expressed by diverse cell populations. We found that among different cell types in the GBM microenvironment, BTICs are the primary source of the ECM molecules. Multiomic analysis showed that biglycan was mostly produced by injury response BTICs which in turn drive their proliferation. We further show biglycan interacted with LRP6 and activated the Wnt/β-catenin pathway in BTICs.

Previous studies suggested elevation of different molecules at the tumor site which mediated tumor cell proliferation. However, there is limited research on the composition of the tumor microenvironment using unbiased analyses. More importantly, cell niche information is lost following tissue dissociation in scRNA-seq studies, limiting its potential to dissect the composition of tumor microenvironment. Spatial transcriptomics allowed us to map the spatial transcriptome profile of the GBM microenvironment in an unbiased manner. Among upregulated genes in the tumor area, we found elevation of 10 ECM molecules including different types of collagens (*Col9a1, Col9a2, Col9a3, Col2a1, Col11a1, Col11a2)*. In line with our findings, GBM tissues have higher levels of collagen expression than normal brain tissues ^20–22^. Furthermore, higher collagen gene expression was associated with a poor prognosis in GBM patients ^23^. We found different cell types in the tumor microenvironment, including tumor cells, oligodendrocytes and immune cells to express collagens. In addition to regulatory functions for tumor cells, collagens also affect the immunosuppressive microenvironment of GBM ^23^. Moreover, collagen genes are involved in the epithelial-mesenchymal transition process of GBM. Collagens were shown to be significantly upregulated in mesenchymal GBM compared to other subtypes ^20, 23^. Interestingly, Col1a1 has been proposed as a therapeutic target for diminishing tumor heterogeneity and mesenchymal phenotype in GBMs ^24^. However, the functions of other collagens members identified in our study in promoting intratumoral heterogeneity are unknown. Another intriguing finding from our research was the increased expression of COL9A3 in developmental than injury response BTICs. The functions of COL9A3 in regulating BTIC heterogeneity and their downstream signaling pathways need more investigation. Fibromodulin (FMOD) was another ECM molecule that was raised in our spatial and single-cell transcriptomic analysis of GBM tissues. Interestingly, fibromodulin through interaction with collagen promotes glioma cell migration and invasion ^25^.

Upregulation of genes encoding CSPGs including versican and CSPG4 was also observed in the GBM microenvironment compared to normal tissues. Other studies, like ours, have shown higher levels of CSPGs in GBM tissues than in normal brains ^26, 27^. Also, expression of CSPGs and enzymes involved in chondroitin sulphate polymerization is elevated in the GBM microenvironment and corresponds with high tumor grade and poor survival ^28–30^. Moreover, CSPGs expression enhances proliferation of tumor cells ^28^. Notably, CSPG4-positive GBM cells overexpress genes associated with aggressive tumorigenicity, such as mitosis and cell cycle module genes ^30^. This is consistent with our findings of elevated CSPG4 expression in tumor area with high proliferative activity. Although versican is produced by both tumor cells and macrophages, we found that CSPG4 is primarily expressed by tumor cells including BTICs. One study has proposed CSPG4 as an antigen target for chimeric antigen receptor (CAR)–redirected T cells in GBM ^31^. CSPG4 is expressed by injury response and developmental BTICs, suggesting its potential for targeting BTICs in GBMs despite BTIC heterogeneity.

We observed significant elevation of biglycan, a proteoglycan family member, in both mouse and human tissues of GBM than healthy brains. Notably, bigycan is mostly expressed by injury response BTICs. Biglycan expression has been shown to be elevated in a variety of solid cancers ^32^. However, little information is available about functions of biglycan in different types of cancer including GBM. The TCGA and scRNA-seq analysis revealed that the mesenchymal subtype of GBM had significantly higher transcript levels of biglycan than the other subtypes. In line with this finding higher correlation was found between biglycan and CD44 (mesenchymal marker) in the culture of human BTICs. Biglycan was discovered to bind with LRP6, hence activating the Wnt/β-catenin pathway and cell growth. Activation of the Wnt/β-catenin pathway is associated with the epithelial-mesenchymal transition ^33^. Consistent with our observation, we found elevated expression of CD44 in mice implanted with *Bgn* overexpressing BTICs. Because the mesenchymal state is linked with a poor prognosis, the biglycan-LRP6 axis may be a new therapeutic target in GBMs.

In summary, comprehensive spatial transcriptomic and scRNA-seq analyses of the mouse and human GBM microenvironment reveal unique transcriptional alterations in the expression of ECM genes associated with the heterogeneity of BTICs in the tumor microenvironment. Exploiting these datasets highlights biglycan which is produced by injury response BTICs and contributes to their proliferation through interaction with LRP6. This study points to biglycan-LRP6 axis as a new therapeutic target to mitigate BTIC proliferation and mesenchymal program in GBMs. New investigations into other gene signatures identified in this study may provide new candidates for treating tumor progression.

## Acknowledgements

We thank the generosity of Anita C. Bellail and Chunhai Hao from Indiana University School of Medicine who kindly provided the human GBM specimens. We acknowledge the Hotchkiss Brain Institute Advanced Microscopy Platform and the Cumming School of Medicine for support and use of Leica TCS SP8 and image analysis platforms. We thank the Snyder Institute’s Live Cell Imaging Resource laboratory at the University of Calgary. We also thank the BTIC Core headed by S. Weiss, G. Cairncross and A. Luchman for isolating BTIC lines from patient-resected specimens and the flow cytometry core, the UCDNA sequencing, and genetic analysis laboratory at the University of Calgary. R.M. was supported by a fellowship from the University of Calgary’s Eyes High program. This study was supported by grants from the Canadian Institutes of Health Research and the Canadian Cancer Society.

## Author Contributions

R.M. conceived the project, designed, performed, analyzed experiments, and wrote the first draft of the paper. C.D.M. prepared the mouse GBM spatial transcriptomic libraries and was key for the spatial transcriptomics and scRNA-seq analysis. A.N and M.G generated and analyzed scATAC-seq datasets. M.K. and P.B. performed data mining of TCGA databases. M.L. helped with analysis of the spatial transcriptomics. V.W.Y. co-conceived the project, provided support and experimental design, supervised the overall study, and critically edited the manuscript. All authors reviewed and edited the manuscript.

## Materials and Methods

### Animal experimental models

All mice used in this study were on C57BL/6 background. Charles River provided six-to 8-week-old female wild-type mice. All experiments were conducted with ethics approval from the Animal Care Committee at the University of Calgary under regulations of the Canadian Council of Animal Care.

### Human tissues

Human GBM tissues for immunohistochemistry (n = 4; Supplementary Data 2) were obtained in compliance with methods authorised by Indiana University’s Institutional Review Board. Fresh tissues were surgically extracted from patients, snap frozen, and kept at 80°C. The tissues were fixed in formalin and histologically processed for paraffin-embedded blocks. The neuropathologist examined the hematoxylin and eosin–stained slides. To collect tumor biopsies, all patients gave written informed permission. AMSBIO provided normal human cerebral cortex for this study (catalog number HP-210).

### Analysis of extracellular matrix (ECM) gene expression in GBMs from TCGA

Gene expression data for TCGA-GBMs was downloaded with the TCGABiolinks and recount2 R packages. Extensive data processing was performed on R. Box-plots associating ECM gene expression with type of sample were generated with the ggbetweenstats function from the ggstatsplot R package by Indrajeet Patil. Multivariate linear model was visualized with a dot and whisker plot with the ggcoefstats function from the ggstatsplot R package. Microarray gene expression data for TCGA-GBMs was downloaded from GDAC Firehose(“GBM.transcriptome ht_hg_u133a broad_mit_edu Level_3 gene_rma data.dat a.txt”). Corresponding clinical data was downloaded from supplementary table 2 by Ceccarelli et. al ^34^. These datasets were imported into R for analysis. Extensive data processing was performed on R. A box-plot associating GBM subtypes with *BGN* expression was generated with the ggstatsplot R package by Indrajeet Patil. For correlation analysis between gene expression, datasets including TCGA GBM RNA-seq and Agilent 4502 were processed via online GBM RNA-seq analysis platform GLIOVIS (http://gliovis.bioinfo.cnio.es/)^35^.

### Generation of GBM patient derived BTICs

Human BTIC lines were generated from resected tissues of GBM patients, as previously reported ^36^. We grew cells in a 5% CO2 incubator in a serum-free NeuroCult NS-A basal medium (STEMCELL Technologies) supplemented with NeuroCult™ Proliferation Supplement (STEMCELL Technologies), 20 ng/mL EGF (PeproTech Inc.), 20 ng/mL FGF (PeproTech Inc.), and 0.5 percent heparin solution (STEMCELL Technologies). BTIC spheres were physically dissociated and seeded onto T-75 culture flasks to proliferate the cells. Within the University of Calgary BTIC Core, these lines were cultured chronologically, preserved, and verified. Identification of the three human BTIC lines (BT048, BT53M and BT073) has been reported previously ^37^.

### Isolation and characterization of mouse BTICs

The mouse BTIC line mBT0309 in the C57BL/6 background was isolated, characterised, and maintained in BTIC medium, as previously reported ^12^. These lines have been demonstrated to mimic critical aspects of a human GBM ^12^.

### Live cell imaging of BTIC growth

For the sphere formation assay, freshly dissociated cells were plated at 10,000 cells per well in 100 μL of BTIC medium into a 96-well flat bottom plate in the presence or absence of 10 μg/mL recombinant human biglycan (R&D Systems, 2667-CM-050). We monitored cell growth using a real-time cell imaging system (IncuCyte live-cell ESSEN BioScience Inc). Images of the cells were taken after 48-72 hours. The resultant number of spheres above the 100-μm-diameter cutoff, a convenient parameter to describe growth characteristics, was analyzed as previously described ^38, 39^.

### Intracranial tumor implantation

After sphere dissociation of firefly luciferase (FL)–expressing BTIC0309, 50,000 viable cells were resuspended in 2 μl of phosphate-buffered saline (PBS) and implanted stereotactically into the right striatum of mice as described previously ^36^. After implantation, we daily monitored animals to assess weight loss and physical/neurological abnormalities.

### In vivo bioluminescence imaging of mice

C57BL/6 mice were intraperitoneally injected with 150 mg/kg of D-Luciferin, Potassium Salt (Gold Biotechnology, LUCK-1G) in DPBS (no calcium or magnesium) 10 minutes before bioluminescence imaging. Mice were anesthetized with isoflurane (2.5% vaporised in O2) and shaved to reduce signal attenuation caused by black hair. Imaging was conducted using Xenogen IVIS 200 system (Xenogen) (auto exposure time). Total photon flux (photons per second) was measured from a defined area of interest using Living Image software for analysis (Xenogen).

### Preparation of libraries for spatial transcriptomics

The Visium Spatial Gene Expression platform (10X Genomics) was used for spatial transcriptomics analysis. Brain tissues were processed according to the manufacturer’s instructions. Briefly, after tumor implantation, when the tumor attained a substantial size as determined by in vivo bioluminescence imaging, brain tissues were extracted and snap frozen using liquid nitrogen before being implanted in FSC 22 Frozen Section Media (Leica). Using a cryostat (ThermoFisher Scientific), brain tissues were cut coronally into 10 μm sections and placed on the capture area of the Visium Spatial Gene Expression Slide. For this study, 2 brain sections from tumor implanted mice and 1 brain section from mice without tumor implantation were used. Tissue optimization was carried out using the Visium Spatial Tissue Optimization Slide and Reagents Kit in accordance with the manufacturer’s instructions. The optimum tissue permeabilization duration was found to be 10 minutes. Brightfield hematoxylin and eosin (H&E) high-resolution images were taken using the EVOS FL Auto Imaging System (ThermoFisher Scientific) using a 10X objective so as to differentiate tumor and non-tumor regions. The subsequent tissue permeabilization, cDNA amplification, barcoding and library construction steps were proceeded with as outlined in the Visium spatial gene expression user guide (CG000239). Prepared libraries were loaded at 300□pM and sequenced on a NovaSeq 6000 System (Illumina) using a NovaSeq 200 Cycle S1 Flow Cell. Sequencing was performed using the following read protocol: read 1: 28 cycles; i7 index read: 10 cycles; i5 index read: 10 cycles; and read 2: 90 cycles.

### Demultiplexing and analysis of spatial transcriptomics

The sequencing depth obtained ranged from approximately 143–183□×□10^6^ reads per library and 45-100,000 mean reads/spot. The base call (BCL) files were processed with the 10x Genomics Space Ranger software v.1.2, which uses STAR v.2.5.1 for genome alignment, against the mm10 mouse reference dataset. The count files generated for each library were then aggregated with normalization set to ‘Mapped’. The aggregated cloupe file was visualized in 10x Genomics Loupe Browser software 5.0. The tumour area was demarcated based on H&E and high expression of proliferation markers. DEG analysis, UMAP plots, violin plots and heat map of DEGs were all performed within Loupe Browser.

### scATAC-seq

Single-cell ATAC-seq analyses were performed on a previously published adult glioblastoma dataset (GSE139136)^8^. Copy number analysis and cycling analysis were performed using Copy-scAT ^40^. Data analysis was performed using Signac ^41^. Datasets were pooled and normalized using mean signal across a randomly selected subset of peaks. ChromVAR was used to infer motif activities ^42^, followed by Non-negative Matrix Factorization (NMF) analysis to decompose cell states within the tumour. Tightly correlated NMF decomposition-based states were pooled to generate two distinct states, and cells with high scores of either state were termed stem or differentiated, respectively, with cells having an intermediate score being termed intermediate. Gene activity at the *BGN* locus was computed using the GeneActivity method in Signac. Comparisons between *BGN* activity in modules were performed using an unpaired T test with Welch’s correction.

### Analysis of public scRNA-seq datasets

The counts matrix provided for the publicly available datasets used was downloaded and converted into a Seurat object using the package Seurat v3 in R ^43^. The data were filtered for the following parameters: cells with >50 genes and the percentage of mitochondrial genes<15%. Data from all 12 libraries were then integrated and normalized with the SCTransform75 wrapper in Seurat using all 17,607 features. A PCA reduction was performed, and 17 significant PCA dimensions were accounted for. Clusters were determined using the FindNeighbours and FindClusters functions, and clustering was performed with a resolution of 0.5. Manual annotation of clusters was performed based on the expression of lineage-specific signature genes. To display cell clusters, RunTSNE with PCA reduction was utilized. DEGs for one cluster (versus all cells in other clusters) was determined through the FindMarkers function. Only DEGs with a statistically significant P value of 0.05 were included for analysis. For each cluster, the DoHeatmap functionality in Seurat was used to plot heat map DEGs of interest. Dot plots depicting the average expression of hallmark genes and genes of interest expressed as a percentage in each of the clusters were created using the DotPlot program. The FeaturePlot and VlnPlot functions were used to create graphs that depict gene expression in cell clusters or sample groups.

### Confocal immunofluorescence microscopy

We evaluated formalin-fixed paraffin-embedded specimens from GBM patients and frozen sections from mice. For formalin-fixed paraffin-embedded specimens, the sections of 10 μm in thickness were subjected to deparaffinization and antigen retrieval using heat-mediated antigen retrieval with tris/EDTA buffer (pH 9). Tissues were blocked by incubation for 1 hour at room temperature with horse-blocking solution [PBS, 10% horse serum, 1% bovine serum albumin (BSA), 0.1% cold fish stain gelation, 0.1% Triton X-100, and 0.05% Tween 20]. Tissues were then incubated overnight at 4°C with following antibodies resuspended in antibody dilution buffer (PBS, 1% BSA, 0.1% cold fish stain gelation, and 0.1% Triton X-100): biglycan (1:500; Novus Biologicals, NBP1-84971), SOX2 (1:200; Thermo Fisher Scientific, 14-9811-82), human Ki67 (1:400; Cell Signaling Technology, 9449), mouse Ki67 (1:500; Abcam, ab15580), β-catenin (1:800; Cell Signaling Technology, 37447S) and CD44 (1:500; Abcam, ab254530). The corresponding fluorophore-conjugated secondary antibodies (1:500; Jackson ImmunoResearch Laboratories or Thermo Fisher Scientific) were then added with 4′,6-diamidino-2-phenylindole (DAPI) (1:1000). Fluoromount-G (SouthernBiotech, 0100-01) was used to adhere coverslips to the slides. Laser confocal immunofluorescence images were obtained at room temperature using the Leica TCS SP8 laser confocal microscope with a 25/0.5 numerical aperture water objective. Samples were excited by 405-, 552-, and 640-nm lasers and detected by one low dark current Hamamatsu photomultiplier tube detectors and two high-sensitivity hybrid detectors on the SP8. The following settings were used to capture images of samples: 8 bits, bidirectional scanning in a z-stack, 4× frame averaging, 1 airy unit pinhole, 0.75 zoom, and 2048 by 2048 pixels x-y resolution. To improve contrast and decrease saturation, the identical laser, gain, and offset settings were utilized for all samples in each series of trials. As a control, a sample slide stained with the secondary antibodies and DAPI was always included in each experiment. Each sample of human and mouse brain slices yielded three and four fields of vision (FOVs). Leica Application Suite X was used to take the images, while Imaris was used to accomplish the picture thresholding and 3D rendering (Bitplane).

### Cell proliferation analysis Click-iT EdU-labeling assay

Cells were treated with 1μM EdU for 24 hours for the Click-iT EdU proliferation assay. Following cell permeabilization, EdU staining was conducted according to the manufacturer’s instructions using the Click-iT EdU Alexa Fluor 488 Flow Cytometry Assay Kit (Thermo Fisher Scientific, C10425). Attune NxT flow cytometry was used for analysis of cells (Thermo Fisher Scientific). FlowJo version 10.7.2 was used to analyse the data (Treestar).

### Confocal image analysis

The z-stack confocal images of spinal cords were analyzed with ImageJ (Fiji, National Institutes of Health). Briefly, maximum intensity projections were generated for each channel/marker and converted from 8-bit to RGB files. Positive signal was determined using the Color Threshold for all markers except DAPI. Threshold was used for DAPI. The Watershed algorithm was used to separate connected cells for quantification of DAPI positive cells. The ‘analyze particles’ function was used to create a mask to quantify the positive signals in each FOV. The same color brightness threshold values, as well as particle analysis size and circularity parameters, were applied consistently across all samples for each experimental set. For representative images shown, maximum intensity projection values of each channel/marker were merged and displayed using pseudo colors. Only the brightness and contrast settings were changed between samples to improve visual presentation.

### Immunofluorescence staining of cells

BTIC was seeded in a 96-well Black/Clear Flat Bottom TC–treated Imaging Microplate (Falcon, 353219) at a density of 10,000 cells per well in BTIC medium. Wells were already coated with 10 g/mL Laminin (Millipore, CC095) to attach BTICs to the plate. After incubation at 37°C, cells were washed with PBS and then fixed in 4% paraformaldehyde for 10 min. Cells were permeabilized using 0.25% Triton X-100 for 5 minutes and then blocked with Intercept® (PBS) Blocking Buffer (LI-CORE, 927-70001) for 1 hour at room temperature. Primary antibodies for biglycan (1:500; Novus Biologicals, NBP1-84971), CD44 (1:500; Abcam, ab254530), β-catenin (1:3000; Cell Signaling Technology, 37447S) diluted in Blocking Buffer were incubated overnight at 4°C. Cells were subsequently incubated with corresponding fluorophore-conjugated secondary antibodies (1:500 Jackson ImmunoResearch Laboratories or Thermo Fisher Scientific) at room temperature for 1 hour and washed, and nuclei were counterstained with Hoechst (1:100; Sigma-Aldrich) and imaged using ImageXpress (Molecular Devices). Multiwavelength cell scoring analysis in the MetaXpress High-Content Image Acquisition and Analysis Software (Molecular Devices) was used to quantify cell numbers. ImageJ was used to merge and display each channel/marker for the sample in pseudo colors for the representative pictures exhibited. Only the brightness and contrast settings were altered, and they were changed consistently between samples to better visual presentation.

### Lentivirus packaging

Lentivirus production was performed using 293FT cells grown to 90% confluency on five, 15 cm^2^ cell culture plates, by cotransfecting 112.5 µg pHAGE PGK-GFP-IRES-LUC-W (a gift from Darrell Kotton (Addgene plasmid #46793;http://n2t.net/addgene:46793; RRID:Addgene_46793)), 73 µg psPAX2 and 39.5 µg pMD2.G (gifts from Didier Trono (Addgene plasmid # 12260; http://n2t.net/addgene:12260; RRID:Addgene_12260)) using CaPO4 precipitation transfection ^44^. Cell culture supernatant was collected and pooled every 12-16 h, centrifuged at 500 x g for 5 min and filtered with a 0.45 µm syringe filter for a total of 3 harvests. The lentivirus-containing supernatant was underlaid with 20% sucrose in PBS and centrifuged at 50,000 x g for 2 h using a Beckman SW28 ultracentrifuge rotor. The supernatant was discarded and the lentivirus pellet was resuspended in sterile PBS and frozen at −80oC in 20 ul aliquots. The lentivirus titer determined using the qPCR lentivirus titer kit was ~108 IU/mL (Applied Biological Materials).

### Plasmid construction

The DNA sequence encoding mouse biglycan (NCBI accession NM_001411776) was PCR amplified from mouse whole brain cDNA whereas the human biglycan (NCBI accession BC002416) sequence was obtained from the The Center for Cancer Systems Biology Human ORFeome collection (Horizon Discovery Biosciences). Both human and mouse *BGN* were subcloned into the PB-CMV-MCS-EF1α-Puro piggybac expression vector (System Biosciences) using the 5’ EcoRI and 3’ BamHI restriction sites. The sequences of all constructs were verified by Sanger DNA sequencing.

### Statistics analysis

Data were collated in Microsoft Excel and graph creation and data analysis were performed in GraphPad Prism 9.4.0. To examine statistically significant differences between the means of two or more treatment groups and the control group, one-way ANOVA with Tukey’s multiple-comparison test was performed. The significance of data with only two groups was tested using a two-tailed, unpaired t-test. As stated in the figure legends, asterisks indicate significance; *P<0.05, **P<0.01 and ***P<0.001. There was no sample size calculation. The sample size was determined using previously published results, the expense of the experiment, the practicality of the experiment, and the availability of sex-and age-matched mice. In violin plots, center lines represent median and two quartile lines. The Pearson method was used in the correlation analysis between gene expressions. The data were all given as mean ±SEM. Unless otherwise specified, all data given are indicative of two to three separate studies with comparable findings. Mycoplasma was commonly detected in cell cultures. Blinding was not conducted.

### Data availability

Spatial transcriptomic datasets reported in this paper are available to download from the NCBI Sequence Read Archive with BioProject (to be completed prior to publication). Additional code is available upon request.

## Supplementary Figures

**Supplementary Figure 1. Quality control analysis of spatial transcriptomics**.

**A-D**. Images show distribution of nCount_spatial (number of transcripts), nFeature-spatial (number of genes), percent_mito (mitochondria content) and percent_hb (hemoglobin content) across tissue section in tumor and normal mouse brains. **E-H**. Violin plots displaying the QC metrics and distribution of cells by the nCount_spatial, by the nFeature-spatial, by the percent_mito and by percent_hb.

**Supplementary Figure 2. Spatial gene expression of ECM molecules in mouse brain tumors**

**A**. Spatial distribution of the 10 ECM genes in tumor 2. **B**. Violin plots comparing the expression levels of ECM genes from two tumor mice compared to one normal mouse. C. Violin plots displaying the expression levels of ECM transcripts amongst the 8 spatial clusters.

**Supplementary Figure 3. Quality control analysis of scRNA-seq. A**. Violin plots show the distribution of cells from scRNAseq of 7 GBM patients by the nFeature_RNA (number of genes detected per cell), by the nCount_RNA (number of UMIs per cell), by the percent_mito (percent mitochondrial content). **B**. Graph shows the top 2000 variable features with the top 10 labeled. **C**. Heat map shows the top 10 differentially expressed genes amongst the 17 clusters. A t-test was used to determine differentially expressed genes, and a comparison was done between cells in each individual cluster and cells in all other clusters. Yellow indicates upregulation, purple indicates downregulation.

**Supplementary Figure 4. Additional analysis showing the correlation between biglycan and mesenchymal phenotype of GBM. A**. Analysis of *BGN* transcript expression in different GBM subtypes via The TCGA database. **B**. Correlation between *BGN* and *CD44* expression in the TCGA-GBM Agilent-4502A (left) and TCGA-GBM RNA-seq (right) database. **C**. Expression of *BGN* in four different GBM cellular states including astrocyte-like (AC-like), mesenchymal-like (MES-like), oligodendroglial progenitor cell-like (OPC-like), and neural progenitor cell-like (NPC-like) signatures (meta-modules) using relative expression of each multi-gene signature (relative meta-module score) previously described (Neftel et al.). **D**. Dot plot showing the expression of various mesenchymal signature genes along with *BGN* gene across the 17 cell clusters identified through scRNA-seq analyzing of 44,712 cells from 7 GBM patients. The size of the dot corresponds to the percentage of cells expressing the gene in each cluster. The color represents the average gene expression level. Dashed lines show clusters with higher expression of *BGN* gene.

**Supplementary Figure 5. Spatial expression of stemness genes in tumor areas. A**. Spatial distribution of stemness genes across tissue section in tumor and normal mouse brains. **B**. Violin plot comparing the expression levels of stemness genes between tumor and normal brains. **C**. Violin plot comparing the expression levels of stemness genes between spatial clusters.

**Supplementary Figure 6. Additional analysis of human GBM specimens for biglycan and BTICs. A, B**. Representative confocal images of human specimens resected from two GBM patients (110907 and 110801). The tissues were labeled with DAPI, SOX2 and biglycan. **C, D**. 3D reconstruction of images of SOX2, and biglycan in human GBM tissues. **E-G**. Representative confocal images of three human normal brains labeled with DAPI, SOX2 and biglycan. **H**. Human GBM tissue was lebeled with secondary antibody as a control.

**Supplementary Figure 7. Expression of biglyan in BTICs in the mouse GBM model. A**. Representative confocal images of brain tissues resected from mice implanted with syngeneic mouse. The tissues were labeled with DAPI, SOX2 and biglycan. **B**. Representative 3D reconstruction of image of SOX2, and biglycan in a mouse GBM tissue. **C**. Representative confocal images of mouse tumor. The tissues were labeled with DAPI, SOX2 and biglycan. **D**. Representative confocal images of normal mouse brain (without tumor implantation). The tissues were labeled with DAPI, SOX2 and biglycan.

**Supplementary Figure 8. Additional analysis of in vitro proliferative effects of biglycan. A**. Bar graphs comparing the ATP proliferation levels between BTICs treated with recombinant biglycan and control cells with no treatment. **B**. Bar graphs comparing the ATP proliferation levels between *BGN* overexpressing and control human BTICs. **C**. Forward scatter vs side scatter-area (FSC-A vs SSC-A) gating was utilised to detect and eliminate debris while identifying cells of interest based on size (FSC-A) and granularity (SSC-A). We removed doublet cells from our study by employing two-step gating of FSC-A versus FSC-height (FSC-H) and SSC-A vs SSC-height (SSC-H). Data are representative of two to three separate experiments. Means were compared between groups by unpaired (two-tailed) t test: *P < 0.05, **P < 0.01, ***P < 0.001.

**Supplementary Data 1** (Microsoft Excel format). Differential gene expression between 8 clusters identified through spatial transcriptomics of brain tissues from GBM mouse model.

**Supplementary Data 2** (Microsoft Excel format). Characterization of patients whose resected tissues were used for confocal microscopy.

